# Association mapping and haplotype analysis of the pre-harvest sprouting resistance locus *Phs-A1* reveals a causal role of *TaMKK3-A* in global germplasm

**DOI:** 10.1101/131201

**Authors:** Oluwaseyi Shorinola, Barbara Balcárková, Jessica Hyles, Josquin F. G. Tibbits, Matthew J. Hayden, Katarina Holušova, Miroslav Valárik, Assaf Distelfeld, Atsushi Torada, Jose M. Barrero, Cristobal Uauy

## Abstract

Pre-harvest sprouting (PHS) is an important cause of quality loss in many cereal crops and is particularly prevalent and damaging in wheat. Resistance to PHS is therefore a valuable target trait in many breeding programmes. The *Phs-A1* locus on wheat chromosome arm 4AL has been consistently shown to account for a significant proportion of natural variation to PHS in diverse mapping populations. However the deployment of sprouting resistance is confounded by the fact that different candidate genes, including the tandem duplicated *Plasma Membrane 19 (PM19)* genes and the *mitogen-activated protein kinase kinase 3 (TaMKK3-A)* gene, have been proposed to underlie *Phs-A1*. To further define the *Phs-A1* locus, we constructed a physical map across this interval in hexaploid and tetraploid wheat. We established close proximity of the proposed candidate genes which are located within a 1.2 Mb interval. An association analysis of diverse germplasm used in previous genetic mapping studies suggests that *TaMKK3-A*, and not *PM19*, is the major gene underlying the *Phs-A1* effect in European, North American, Australian and Asian germplasm. We identified the non-dormant *TaMKK3-A* allele at low frequencies within the A-genome diploid progenitor *Triticum urartu* genepool, and show an increase in the allele frequency in modern varieties. In UK varieties, the frequency of the dormant *TaMKK3-A* allele was significantly higher in bread-making quality varieties compared to feed and biscuit-making cultivars. Analysis of exome capture data from 58 diverse hexaploid wheat accessions identified fourteen haplotypes across the extended *Phs-A1* locus and four haplotypes for *TaMKK3-A*. Analysis of these haplotypes in a collection of UK and Australian cultivars revealed distinct major dormant and non-dormant *Phs-A1* haplotypes in each country, which were either rare or absent in the opposing germplasm set. The diagnostic markers and haplotype information reported in the study will help inform the choice of germplasm and breeding strategies for the deployment of *Phs-A1* resistance into breeding germplasm.

## Introduction

Pre-harvest sprouting (PHS) refers to the too-early germination of physiologically matured grains while still on the ear, but before harvest. PHS is primarily caused by insufficient levels, or rapid loss, of seed dormancy and is an important cause of quality loss in many cereal crops (Li et al., 2004; Fang and Chu, 2008). This is particularly relevant in wheat due to its detrimental effects on bread-making potential which represents the most common use of wheat grains globally (Simsek et al., 2014). PHS is believed to be a modern phenomenon, as progenitor and wild wheat species generally display high levels of seed dormancy (Gatford et al., 2002; Lan et al., 2005). Selection for reduced seed dormancy during domestication and modern breeding programmes allowed for more uniform seed germination and rapid crop establisment (Nave et al., 2016). However, this also resulted in higher level of susceptiblity to PHS in modern wheat varieties (Barrero et al., 2010). In addition to its detrimental effect on quality, PHS also reduces yield and affects seed viability, making resistance to PHS a high priority in many breeding programmes.

Occurrence of PHS is heavily influenced by the environment. PHS is prevalent in wheat growing regions with high levels of rainfall during the period of grain maturation and after-ripening. Increased ambient temperature during this period can further increase the susceptibility of grains to sprouting (Barnard and Smith, 2009; Mares and Mrva, 2014). This enviromental dependency of PHS constitutes a constraint in selecting for PHS resistance in field conditions. In addition, resistance to PHS is highly quantitative and is controlled by numerous quantitative trait loci (QTL) located on all 21 chromosomes of bread wheat (Flintham et al., 2002; Kulwal et al., 2005; Mori et al., 2005; Kottearachchi et al., 2006; Ogbonnaya et al., 2007; Liu et al., 2008; Torada et al., 2008; Xiao-bo et al., 2008; Mohan et al., 2009; Munkvold et al., 2009; Knox et al., 2012; Kulwal et al., 2012; Gao et al., 2013; Lohwasser et al., 2013; Mares and Mrva, 2014; Kumar et al., 2015). This makes resistance to PHS one of the most multi-genic traits in wheat and further highlights the complexity in breeding for this trait.

Despite the multi-genic control of PHS resistance, a locus on chromosome arm 4AL, designated as *Phs-A1*, has been consistently shown to account for a significant proportion of natural variation to sprouting in diverse mapping populations. The *Phs-A1* effect has been identified in at least eleven bi-parental and multi-parent mapping populations derived from diverse germplasm from Australia, UK, Japan, China, Mexico, Canada and Europe (Torada et al., 2005; Ogbonnaya et al., 2007; Chen et al., 2008; Torada et al., 2008; Cabral et al., 2014; Albrecht et al., 2015; Barrero et al., 2015). Physiological evaluation of *Phs-A1* shows that it delays the rate of dormancy loss during seed after-ripening when plants are grown across a wide range of temperatures (13 °C – 22 °C; Shorinola et al., 2016).

Recently, two independent studies by Barrero et al. (2015) and Torada et al. (2016) identified the tandem duplicated *Plasma Membrane 19 (PM19-A1* and *PM19-A2*) genes and a *mitogen-activated protein kinase kinase 3 (TaMKK3-A)* gene, respectively, as candidates for *Phs-A1*. The *PM19* genes were identified through a combined genetic approach using multi-parent mapping populations and transcriptomic analysis of near-isogenic recombinant inbred lines. The *TaMKK3-A* gene was identified through a more traditional positional cloning strategy using bi-parental mapping populations. Each study confirmed the effect of the gene(s) on dormancy through either down-regulation of transcript levels through RNA interference *(PM19)* or transgenic complementation of the susceptible parent with the resistant allele *(TaMKK3-A)*.

It is presently unclear whether the sprouting variation associated with *Phs-A1* across diverse germplasm is due to allelic variation at *PM19* or *TaMKK3-A* alone, or if it’s due to a combination of both genes (Torada et al 2016). Fine-mapping studies (Shorinola et al., 2016) defined *Phs-A1* to a genetic interval distal to *PM19* for UK germplasm, consistent with the position of *TaMKK3-A*. However, a comprehensive understanding of *Phs-A1* diversity taking into account both *PM19* and *TaMKK3-A* genes across a wider set of germplasm is lacking.

In this study, we characterised the *Phs-A1* physical interval in both hexaploid and tetraploid emmer wheat to establish the physical proximity of *PM19* and *TaMKK3-A*. We developed markers for the candidate genes, and showed *TaMKK3-A* alleles to be diagnostic for sprouting resistance in a panel of parental lines from mapping populations in which *Phs-A1* was identified. We used diploid, tetraploid and hexaploid accessions to further trace the origin of the sprouting susceptible *TaMKK3-A* allele and used exome capture data from the wheat HapMap panel (Jordan et al., 2015) to examine the haplotype variation across the *Phs-A1* locus.

## Materials and Methods

### Physical Map Sequence Assembly and Annotation

A fingerprinted Bacterial Artificial Chromosome (BAC) library of flow-sorted 4A chromosome was used for constructing the Chinese Spring *Phs-A1* physical map (accessible at https://urgi.versailles.inra.fr/gb2/gbrowse/wheat_phys_4AL_v2/). Using the high-throughput BAC screening approach described by Cvikova et al. (2015), a sequence database made from three-dimensional pool of BAC clones comprising the Minimum Tilling Path (MTP) was searched for the sequences of *PM19-A1* and *TaMKK3-A*. This identified two positive clones for *PM19-A1* (TaaCsp4AL037H11 and TaaCsp4AL172K12) and three positive clones for *TaMKK3-A* (TaaCsp4AL032F12, TaaCsp4AL012P14 and TaaCsp4AL002F16; Table S1). Using Linear Topology Contig (LTC; Frenkel et al., 2010) BAC clustering information for this library, we identified the BAC clusters (defined as a network of overlapping BACs forming a contiguous sequence) to which these BACs belong. The *PM19-A1*-containing BACs belong to BAC Cluster 16421 which has 20 BACs in its MTP while the TaMKK3-A-containing BACs belong to BAC Cluster 285 comprised of four MTP BACs (Table S1).

DNA of the BACs was extracted using the Qiagen Plasmid Midi Kit (Qiagen, Cat. No. 12143). Eleven of the 20 MTP BACs of Cluster 16421 and the four BACs of Cluster 285 MTP were sequenced on the Illumina MiSeq with 250 bp paired-end reads. An average of 2,105,488 and 2,752,220 paired-end reads per BAC were produced for Cluster 16421 and 285 BACs, respectively. Illumina reads for each BAC were separately assembled using the CLC Bio genomic software (www.clcbio.com). Before assembly, reads were filtered to remove contaminant sequences by mapping to the BAC vector (pIndigoBAC-5) sequence and the *Escherichia coli* genome. *De novo* assembly of reads after contaminant removal was done with the following assembly parameters: Word size: 64 bp; Bubble size: 250 bp; Mismatch cost: 2; Insertion cost: 3; Deletion cost 3; Length fraction: 90%; Similarity fraction: 95%.

The assembled contigs were repeat-masked by BLASTn analysis against the Triticeae Repeat Element Database (TREP: wheat.pw.usda.gov/ITMI/Repeats; Wicker et al., 2000). Gene annotation was performed using the wheat gene models described by Krasileva et al. (2013) and by BLASTX analysis to NBCI nr (blast.ncbi.nlm.nih.gov/Blast). Gene models were also obtained by *ab-initio* gene prediction with FGENESH (Solovyev et al., 2006). Only FGENESH gene models with protein sequence support from NCBI or *Ensembl* Plant protein databases (plants.ensembl.org) were used. Gene models with greater than 90% protein or nucleotide sequence identity and more than 75% sequence coverage to already annotated genes on NCBI or *Ensembl* databases were considered as high confidence genes. Gene models that did not meet these criteria were considered as low confidence genes, and were not analysed further.

#### TaMKK3-A genotyping

A Kompetitive Allelic Specific PCR (KASP; Smith and Maughan, 2015) assay was developed for genotyping the C to A (C>A) causal *TaMKK3-A* mutation reported by Torada et al. (2016). For this, two allele-specific reverse primers *(TaMKK3-A-snp1-res:* TTTTTGCTTCGCCCTTAAGG and *TaMKK3-A-snpA1-sus:* TTTTTGCTTCGCCCTTAAGT) each containing the allele-specific SNP at the 3’ end, were used in combination with a common A-genome specific forward primer (GCATAGAGATCTAAAGCCAGCA). To distinguish the amplification signal produced from each allele specific primer, FAM and HEX fluorescence dye probes (Ramirez-Gonzalez et al., 2015) were added to the 5’ end of *TaMKK3-A-snpA1-res* and *TaMKK3-A-snpA1-sus*, respectively. KASP assays were performed as previously described (Shorinola et al., 2016).

In addition to the KASP assay, a genome-specific Cleavage Amplified Polymorphism Sequence (CAPS) assay (Konieczny and Ausubel, 1993), designated as *TaMKK3-A-caps*, was developed. This CAPS marker is associated with the presence/absence of an *Hpy* 166II restriction site which co-localises with the C>A causal polymorphism in the fourth exon of *TaMKK3-A*. Genome-specific primer pairs (Forward: CACCAAAGAATAGAAATGCTCTCT and Reverse: AGGAGTAGTTCTCATTGCGG) were designed to amplify an 887-bp sequence including the fourth exon. PCR was performed with Phusion High Fidelity polymerase (NEB, UK; Cat No: M0530S) in a 50 μL volume containing 20 % buffer, 0.2 mM of dNTP, 5 μM each of *TaMKK3-A-cap* forward and reverse primers, 3 % of DMSO, 200 - 400 ng of genomic DNA and 0.5 unit of Phusion polymerase (NEB, UK; Cat No: M0530S). Thermal cycling was done with Eppendorf Mastercycler^®^ Pro Thermal Cyclers with the following programme: initial denaturation at 98 °C for 2 mins; 35 cycles of denaturation at 98 °C for 30 s; Annealing at 62 °C for 30 s and extension at 72 °C for 60 s; final extension at 72 °C for 10 mins. Following PCR amplification, a 25 μL restriction digest reaction containing 21.5 μL of the final PCR reaction, 2.5 μL of CutSmart^®^ Buffer (NEB, UK; Cat No: B7204S) and 10 units of *Hpy*166II was incubated at 37 °C for 1 hr. Digest products were separated on a 1.5 % agarose gel.

#### PM19-A1 genotyping

Detection of an 18 bp deletion on the promoter region of *PM19-A1* was carried out using primers TaPM19-A1-5F (GAAACAGCTACCGTGTAAAGC) and TaPM19-A1-5R (TGGTGAAGTGGAGTGTAGTGG) reported by Barrero et al. (2015). PCR reaction mixture contained template DNA, 2.5 mM MgCl**2**, 1.5 mM dNTP, 1.5 μM of each primer, and 1 unit of *Taq* polymerase (NEB). The reaction mixture was made up to a total volume of 10 μl. The PCR conditions were as follows: 3 min at 94°C, followed by 30 cycles of 40 s at 94°C, 40 s at 60°C, and 1 min at 72°C. The last step was incubation for 7 min at 72°C. The PCR products were resolved on a 4% agarose gel and visualized with SYBR green I (Cambrex Bio Science, Rockland, ME.).

### Germplasm for Association Mapping

We genotyped *PM19* and *TaMKK3-A* across 23 wheat varieties previously reported to segregate for *Phs-A1*, including UK (Alchemy, Robigus, Option, Claire, Boxer, Soleil), Australian (Yitpi, Baxter, Chara, Westonia, Cranbrook, Aus1408, Janz, Cunningham, Halberd), Japenese (Kitamoe, Haruyokoi, OS21-5), Mexican/CIMMYT (W7984, Opata M85), Canadian (Leader), Chinese (SW95-50213) and Swiss varieties (Münstertaler). We also genotyped *TaMKK3-A* in accessions from progenitor species *T. urartu* (41 accessions; A^u^ genome), *T. turgidum* ssp. *dicoccoides* (151 accessions; AABB genomes), 804 hexaploid accessions from the Watkins landrace collection (Wingen et al., 2014), and 457 modern European bread wheat varieties from the Gediflux collection released between 1945 and 2000 (Reeves et al., 2004).

### Variant Calling and Haplotype Analysis

We examined the haplotype structure around the *Phs-A1* locus in three different germplasm sets. These included 457 varieties in the Gediflux collection, a panel of 200 Australian varieties, and the wheat Haplotype Map (HapMap) panel consisting of 62 diverse global accessions (Jordan et al., 2015). For the HapMap panel, we selected polymorphic sites as follows. We extracted SNP information from published variant call files (VCF) produced from whole exome capture (WEC) resequencing dataset of the 62 HapMap lines (accessible at www.wheat-urgi.versailles.inra.fr/Seq-Repository/Variations). For this, the corresponding IWGSC contig information for genes represented in the *Phs-A1* physical map were first obtained and used to filter the HapMap VCF for SNP sites located within these contigs. We kept SNP sites with allele frequencies of >5 % and accessions with >80% homozygous calls across SNPs. Allele information at the selected SNP loci was reconstructed for each line using the reference, alternate and genotype field information obtained from the VCF. Haplotype cluster analysis was done with Network 5.0.0.0 (Fluxus Technology Limited, UK) using the Median Joining Network Algorithm.

### Pedigree Visualisation

Pedigree information was obtained from the Genetic Resources Information System for Wheat and Triticale (GRIS, http://wheatpedigree.net/) and the International Crop Information System (ICIS: www.icis.cgiar.org). Pedigree visualisation was performed with Helium (Shaw et al., 2014). The coefficient of parentage (COP) analysis (i.e. the probability that alleles of two individuals are identical by descent) was calculated for all pairwise comparisons of lines within the most prevalent haplotypes (Australian: H1/H2 and H5/H7; UK: H3 and H12). For accuracy, landraces or cultivars with unknown or ambiguous pedigrees were not included in the COP analysis. Diversity within haplotype groups was estimated by the mean calculation of all COPs within each matrix.

## Results

### *TaMKK3-A* and *PM19* are located within a 1.2 Mb physical interval

We constructed an extended physical map across the *Phs-A1* interval to investigate the physical proximity between the *TaMKK3-A* and *PM19* candidate genes. Using *PM19-A1* and *TaMKK3-A* sequences as queries, we screened *in silico* a BAC library of flow-sorted 4AL chromosome arm of the bread wheat cultivar Chinese Spring (CS). *PM19-A1* and *TaMKK3-A* were found on two independent non-overlapping BAC clone clusters which were anchored on the high resolution radiation hybrid map of chromosome 4A (Balcárková et al., 2016). The MTP of Cluster 16421 *(PM19)* was comprised of eleven BAC clones whereas the MTP of Cluster 285 *(TaMKK3-A)* included four BAC clones (Figure 1A; Table S1).

**Figure 1:**
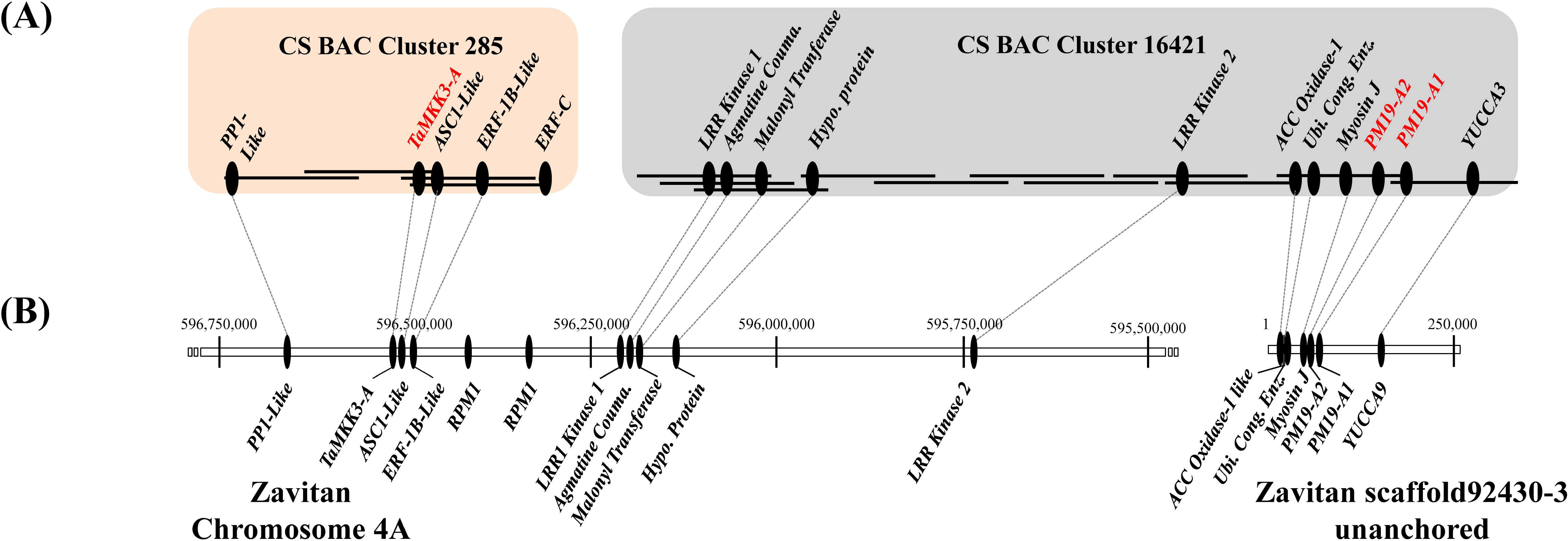
Physical map of the *Phs1-A1* interval in bread wheat Chinese Spring (CS) and wild emmer (Zavitan). (A) *Phs1-A1* interval in CS is covered by two non-overlapping BAC clusters: Cluster 285 (4 BACs) and Cluster 16421 (11 BACs). BACs are represented by solid lines while genes found on the BAC are represented by filled ovals. The proposed candidate genes for *Phs-A1* are in red font. (B) Whole genome assembly of Zavitan wild emmer across the *Phs1-A1* interval. Genes present in both assemblies are joined by dotted lines.

Individual BACs were sequenced, assembled, repeat-masked and annotated for coding sequences. Cluster 16421 included nine high-confidence genes in addition to the *PM19-A1* and *PM19-A2* genes. These included *YUCCA3-like, Myosin-J Heavy Chain protein, Ubiquitin Conjugating Enzyme, Amino-Cyclopropane Carboxylate Oxidase 1 like (ACC Oxidase-1)*, two *Leucine-Rich Repeat Kinases (LRR kinase 1 and LRR kinase 2), Agmatine Coumaroyl Transferase, Malonyl Coenzyme A:anthocyanin 3-O-glucoside-6″-O-malonyltransferase* and a gene encoding for a hypothetical protein. In addition to *TaMKK3-A*, Cluster 285 contained four additional genes including *Protein Phosphatase1-Like (PP1-Like), Activating Signal Co-integrator 1-Like (ASC1-Like), Ethylene Responsive Factor-1B-Like (ERF-1B-Like)* and a gene fragment showing high sequence similarity to *ERF-1B-Like* and as such designated as *ERF-C*. Together, this highlights the presence of at least 16 protein-coding genes across the *Phs-A1* interval in hexaploid bread wheat (Figure 1).

We also characterised the *Phs-A1* interval in the recently constructed assembly of a wild emmer wheat, Zavitan (Figure 1B; Hen-Avivi et al., 2016). This allowed comparative analysis of the *Phs-A1* interval in tetraploid and hexaploid wheat species. Fifteen of the 16 genes found in the CSS physical map were located on two Zavitan scaffolds. Nine of these 15 genes were positioned across a 0.93 Mb interval on the Zavitan 4A pseudomolecule. These included 4 genes from BAC Cluster 285 and five genes from BAC Cluster 16421 (Figure 1). The remaining six genes spanned a 0.13 Mb interval on an unanchored scaffold. On average, the coding sequence identity between CS and Zavitan was 99.7% across the genes shared by both assemblies. We could not find sequence for *ERF-C* in the Zavitan assembly at similar identity. We annotated two genes encoding for disease resistance protein *RPM1* in the Zavitan sequence corresponding to the gap between the two CS BAC clusters. Combining the CS and Zavitan physical maps, the physical region between *TaMKK3-A* and the *PM19* genes was covered and estimated to be approximately 1.2 Mb (Figure 1).

### *TaMKK3-A* is most closely associated with *Phs-A1*

Torada et al. (2016) reported a C>A mutation in position 660 of the *TaMKK3-A* coding sequence (C660A) as being causative of the *Phs-A1* effect. Using alignments of the three wheat genomes, we developed a genome-specific and co-dominant KASP assay for this SNP designated as *TaMKK3-A-snp1*. The *TaMKK3-A-snp1* assay is co-dominant as it distinguished between heterozygotes and homozygotes F_2_ progenies in the Alchemy x Robigus population previously reported to segregate for *Phs-A1* (Shorinola et al., 2016; Figure 2A). We also developed a CAPS marker (Konieczny and Ausubel, 1993) for *TaMKK3-A* to enable genotyping of *Phs-A1* using a gel-based assay. This marker, designated as *TaMKK3-A-cap*, amplifies a genome-specific 887 bp region and is designed to discriminate for the presence of an *Hpy*166II site (GTNNA<u>C</u>) which is lost by the C660A mutation. Dormant lines with the C allele maintain the Hpy166II site which leads to digestion of the 887 bp amplicon into fragments of 605 and 282 bp (Figure 2B). Conversely, non-dormant lines with the A allele lose the *Hpy*166II site and hence remain intact (887 bp) after digestion. As with the KASP assay, the CAPS marker was co-dominant when used to genotype F_2_ progenies (Figure 2B).

**Figure 2:**
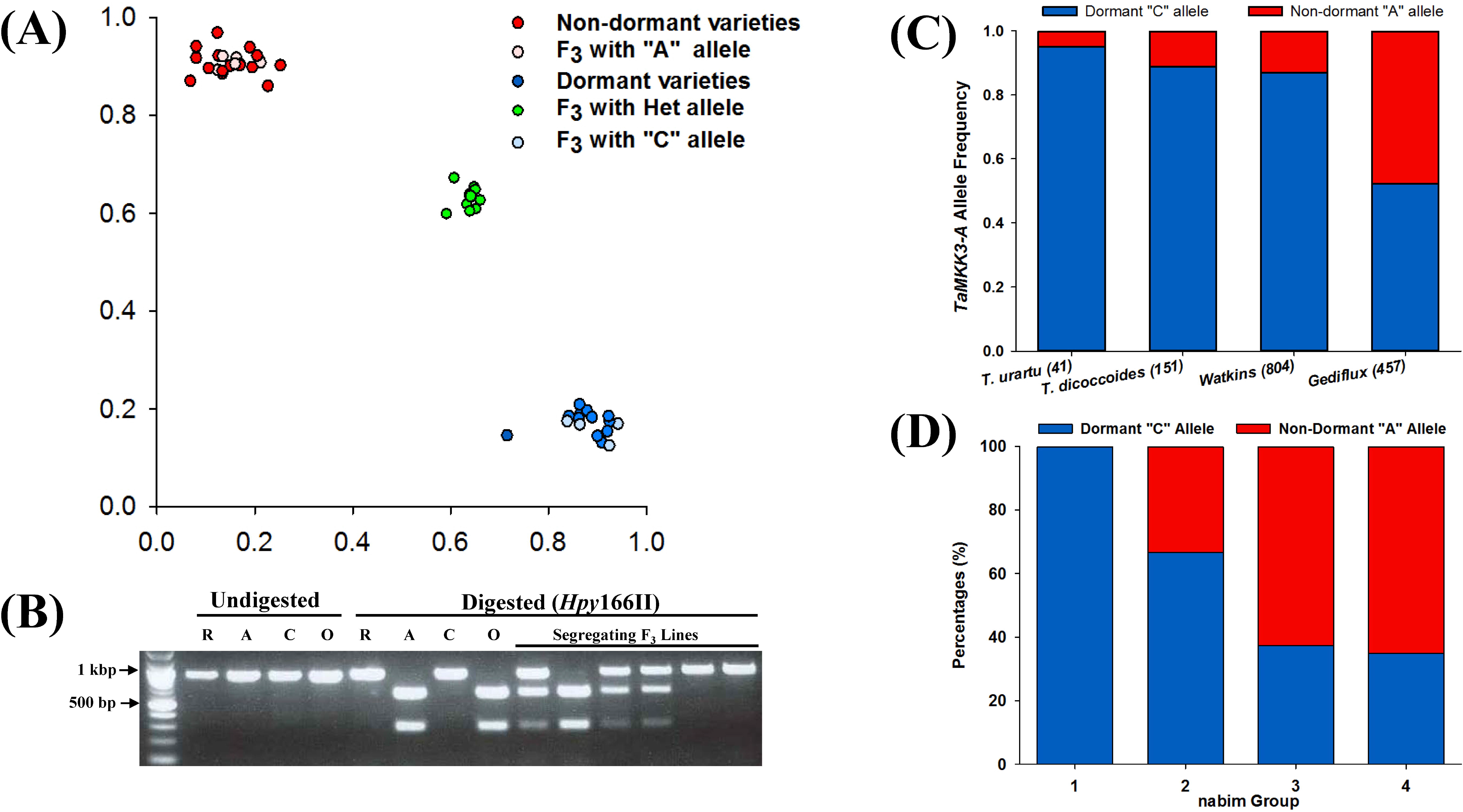
Marker development and allele distribution of *MKK3-A* in ancestral and historic germplasm. (A) Genotype plot of varieties and a F_3_ population segregating for *Phs-A1* using the *TaMKK3-A* KASP assay. (B) Development of co-dominant CAPS marker based on *Hpy*166II restriction digest of the C660A SNP. Non-dormant varieties Robigus (R) and Clare (C); dormant varieties Alchemy (A) and Option (O). (C) Allele frequency of the causal C660A SNP in *T. urartu* and *T. turgidum* ssp. *dicoccoides* accessions and the Watkins and Gediflux collections. The number of lines genotyped in each germplasm set is in parenthesis. (D) *TaMKK3-A* allele distribution in the four wheat end-use groups (nabim 1-4) in the UK.

Using the KASP assay, we genotyped an association panel comprised of the parents of 11 bi-parental mapping populations and a MAGIC population in which *Phs-A1* had previously been reported (Table 1). The *TaMKK3-A-snp1* was polymorphic and diagnostic for *Phs-A1* in all parental lines. Consistent with Torada et al. (2016), non-dormant sprouting-susceptible parents carry the *TaMKK3-A* “A” allele while all the dormant sprouting-resistant parents carry the *TaMKK3-A-snp1* “C” allele (Table 1). We genotyped the same panel for the promoter deletion in *PM19-A1* previously proposed to be causal of PHS susceptibility (Barrero et al., 2015). We found the *PM19-A1* deletion to be linked with the non-dormant *TaMKK3-A* A allele in most, but not all, of these populations. The putative linkage was broken in the dormant Kitamoe, OS21-5 and SW95-50213 parents, whose dormancy phenotypes are not consistent with their *PM19-A1* promoter deletion status, but can be explained by their *TaMKK3-A* genotype (Table 1). This association was confirmed genetically in the SW95-50213 x AUS1408 cross. This population, which did not segregate for the dormancy phenotype in the original work by Mares et al. (2005), is monomorphic for the dormant C allele at *TaMKK3-A*, but segregates for the *PM19-A1* deletion. Similarly, parents of the two populations OS21-5 x Haruyokoi and Kitamoe x Münstertaler segregating for the dormancy phenotype in the work by Torada et al. (2005) are monophormic for the *PM19-A1* deletions, but segregate accordingly for the *TaMKK3-A* causal mutation. These results strongly support *TaMKK3-A* as the most likely causal gene for *Phs-A1* across this highly-informative panel.

**Table 1:** *TaMKK3-A* and *PM19* alleles in *Phs-A1* association panel

| Population | Variety | Origin | Status | *TaMKK3-A* Allele | *PM19-A1* promoter InDel | Reference |
| --- | --- | --- | --- | --- | --- | --- |
| Alchemy x Robigus | Alchemy | UK | Dormant | C | Insertion | Shorinola et al (2016) |
|  | Robigus | UK | Non-dormant | A | Deletion |  |
| Option x Claire | Option | UK | Dormant | C | Insertion | Shorinola et al (2016) |
|  | Claire | UK | Non-dormant | A | Deletion |  |
| MAGIC Population | Yitpi | AUS | Dormant | C | Insertion | Barrero et al (2015) |
|  | Baxter | AUS | Non-dormant | A | Deletion |  |
|  | Chara | AUS | Non-dormant | A | Deletion |  |
|  | Westonia | AUS | Non-dormant | A | Deletion |  |
| Opata x W7984 | W7984 | MEX | Dormant | C | Insertion | Lohwasser et al, (2013) |
|  | Opata | MEX | Non-dormant | A | Deletion |  |
| OS21-5 x Haruyokoi | OS21-5 | JPN | Dormant | C | Deletion | Torada et al, (2008) |
|  | Haruyokoi | JPN | Non-dormant | A | Deletion |  |
| Leader x Haruyokoi | Leader | CAN | Dormant | C | Insertion | Torada et al, (2008) |
|  | Haruyokoi | JPN | Non-dormant | A | Deletion |  |
| Kitamoe x Münstertaler | Kitamoe | JPN | Dormant | C | Deletion | Torada et al, (2005) |
|  | Münstertaler | SUI | Non-dormant | A | Deletion |  |
| Cranbrook x Halberd | Halberd | AUS | Dormant | C | Insertion | Mare et al (2005), Zhang et al (2008) |
|  | Cranbrook | AUS | Non-dormant | A | Deletion |  |
| Janz x AUS1408 | Aus1408 | AUS, SA | Dormant | C | Insertion | Mare et al (2005); Ogbonnaya et al (2007) |
|  | Janz | AUS | Non-dormant | A | Deletion |  |
| SW95-50213 x Cunningham | SW95-50213 | CHN | Dormant | C | Deletion | Mares et al (2005) |
|  | Cunningham | AUS | Non-dormant | A | Deletion |  |
| SW95-50213 x AUS1408* | SW95-50213 | CHN | Dormant | C | Deletion | Mares et al (2005) |
|  | Aus1408 | AUS, SA | Dormant | C | Insertion |  |
| Boxer x Soleil | Soleil | UK | Dormant | C | Insertion | Flintham (2000) |
|  | Boxer | UK | Non-dormant | A | Deletion |  |

\**Phs-A1* was not detected in this population as the DH lines were generally dormant. However, limited number of lines showed transgressive segregation relative to the dormant phenotypes of the two parents.

### Origin and distribution of the *TaMKK3-A* alleles in ancestral and modern germplasm

To examine the origin, distribution and allele frequencies of the causative *TaMKK3-A* C660A SNP, we genotyped a set of 41 *T. urartu* (diploid: AA genome) and 151 *T. turgidum* ssp. *dicoccoides* (tetraploid: AABB genome) accessions. These represent the diploid and tetraploid progenitors of the modern bread wheat A genome on which *Phs-A1* is located. Torada et al. (2016) previously suggested that the non-dormant A allele was the mutant form since the dormant C SNP was conserved across different species. Across *T. urartu* accessions, the C allele was predominant (39 accessions) while the non-dormant A allele was present in only two accessions (5% allele frequency; Figure 2C). Similarly, across *T. dicoccoides* accessions, the dormant C allele frequency was found in 134 accessions while the non-dormant allele was found in 17 accessions (11% allele frequency; Figure 2C). Our results are consistent with Torada et al. (2016) in that the non-dormant A allele is derived from the wild type C allele. In addition, the presence of the A allele across both progenitor species suggests that the mutation predates the hybridization and domestication events that gave rise to modern bread wheat.

We also genotyped the Watkins Collection representing a set of global bread wheat landraces collected in the 1920 and 1930s (Wingen et al., 2014), as well as the Gediflux collection comprised of modern European bread wheat varieties released between 1945 and 2000 (Reeves et al., 2004). The allele frequency of the non-dormant A allele was 13% in the Watkins landrace collection (Figure 2C; Table S2), comparable to that in *T. dicoccoides* (11%). However, the non-dormant A allele frequency in the Gediflux collection was 48% across 457 varieties (Figure 2C). This represents a marked increase of the non-dormant allele in the more modern European collection when compared to European accessions within the Watkins landraces in which the non-dormant A allele was found at a 15% frequency (Table S2).

To determine if the *TaMKK3* dormant allele was associated with improved end-use quality, we genotyped 41 UK varieties representing the four UK market classes (Figure 2D, nabim groups 1-4; nabim, 2014). Of the 13 bread-making quality varieties (groups 1 and 2), eleven (85%) had the dormant *TaMKK3* allele. This frequency was significantly higher (Contingency table χ^2^ = 8.497; *P* < 0.01) than in the 28 biscuit and animal feed varieties (groups 3 and 4) in which the *TaMKK3* dormant allele was only present in 10 varieties (36%).

### *Phs-A1* haplotypes in global germplasm

We next examined the allelic diversity across the extended *Phs-A1* interval (including *TaMKK3-A* and *PM19)* with the aim of elucidating the haplotype structure across this region. For this, we used the SNP Haplotype Map (HapMap) dataset obtained from whole exome capture resequencing of 62 diverse germplasm (Jordan et al., 2015). From this SNP dataset, we obtained data for eight of the sixteen genes found in the *Phs-A1* interval *(PP1-like, TaMKK3-A, ASC1-like, ERF-C, LRR Kinase 1, LRR Kinase 2, PM19-A2* and *PM19-A1)* corresponding to 51 SNPs. To improve the accuracy of the haplotype analysis, we selected accessions with >80% homozygous calls across the selected genes and SNPs with >5% allele frequency. This filtering resulted in 39 SNPs across the eight genes in 58 accessions.

Across the *Phs-A1* interval *(PP1-like* to *PM19-A1)* we identified 14 distinct haplotypes (H1–14; Figure 3A). Haplotypes were comprised of a mix of cultivars, landrace, breeding lines and synthetic population in varying proportion (Figure 3B; Table S3). H1 represented the major haplotype present in 33% of all accessions examined, whereas five haplotypes were relatively infrequent (<5%; H2, H5, H6, H9, H13). Also, we observed haplotype linkage from the *TaMKK3-A* to *LRR Kinase 2* in 76% of the accessions, highlighting possible evidence for limited recombination in this 780 kb interval in global germplasm. Similar haplotype linkage was observed at the tandem *PM19* loci in all but one of the 58 lines.

**Figure 3:**
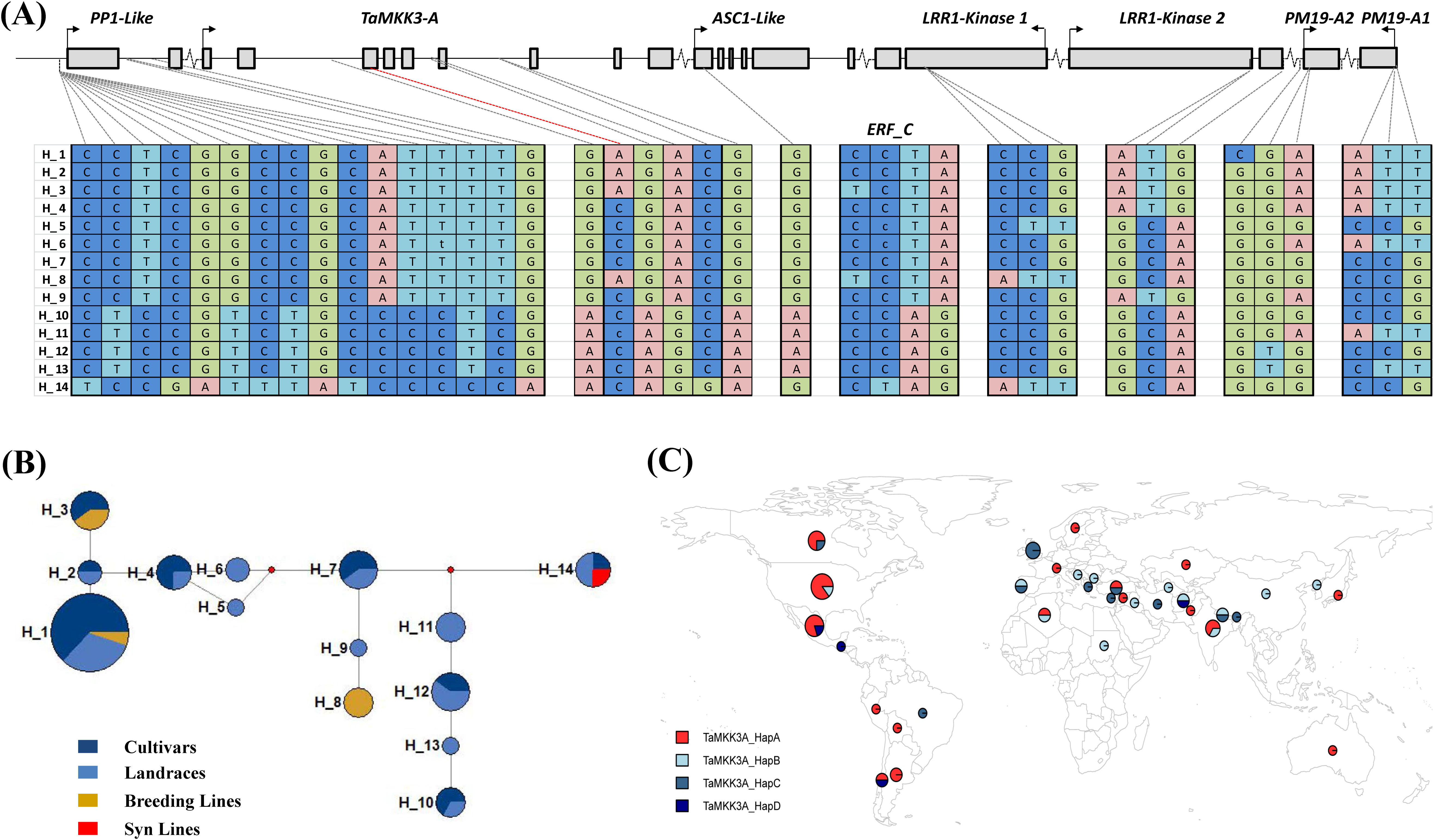
*Phs-A1* haplotype analysis. (A) Structure of 14 haplotypes identified in the HapMap population across 39 SNP loci in the *Phs-A1* interval. SNP loci are ordered based on their physical position on the Zavitan assembly. Exons, intron and intergenic regions are represented by filled boxes, solid lines and breaks, respectively. Note that *ERF-C* is not on the Zavitan assembly. (B) Haplotype network of the 14 haplotypes. The size of each circle corresponds to the number of lines in each haplotype. Blue, light blue, amber and red represent cultivars, landraces, breeding and synthetic line in each haplotype, respectively. (C) Geographical distribution of the four *TaMKK3-A* haplotypes. The size of the circle represents the sample size obtained within each country while each section represents the proportion of the country sample size with the specified haplotype.

Five of the selected SNPs were found in *TaMKK3-A* including the causal C660A SNP in the fourth exon and four additional intron SNPs. These five SNPs defined four distinct *TaMKK3-A* haplotypes (Figure 3C, TaMKK3-A_HapA - D) in the HapMap collection with only one having the non-dormant A allele (TaMKK3-A_HapA). The non-dormant A allele was present in 50% of the HapMap population, consistent with the Gediflux collection (48%).

### Haplotype structure at the *Phs-A1* interval in UK and Australian germplasm

To characterise a larger set of European (Gediflux) and Australian germplasm, we selected seven informative polymorphisms across seven genes from the HapMap dataset and developed KASP assays for these (Table S4). Using these seven assays, we defined 16 haplotypes in the European Gediflux collection (Table S5). This included eleven haplotypes previously identified in the global HapMap dataset and five haplotypes unique to this European germplasm set, although these were relatively infrequent (Figure 4). The UK subpopulation within the Gediflux collection comprised of 176 varieties and contained 11 of the 15 haplotypes identified. Six haplotypes include the dormant *TaMKK3-A* C allele (63% of UK varieties), with the majority of these varieties sharing haplotype H12 (89 of 110 varieties), consistent with the wider Gediflux population (Figure 4). This suggests one main source of PHS resistance in UK and European germplasm.

**Figure 4:**
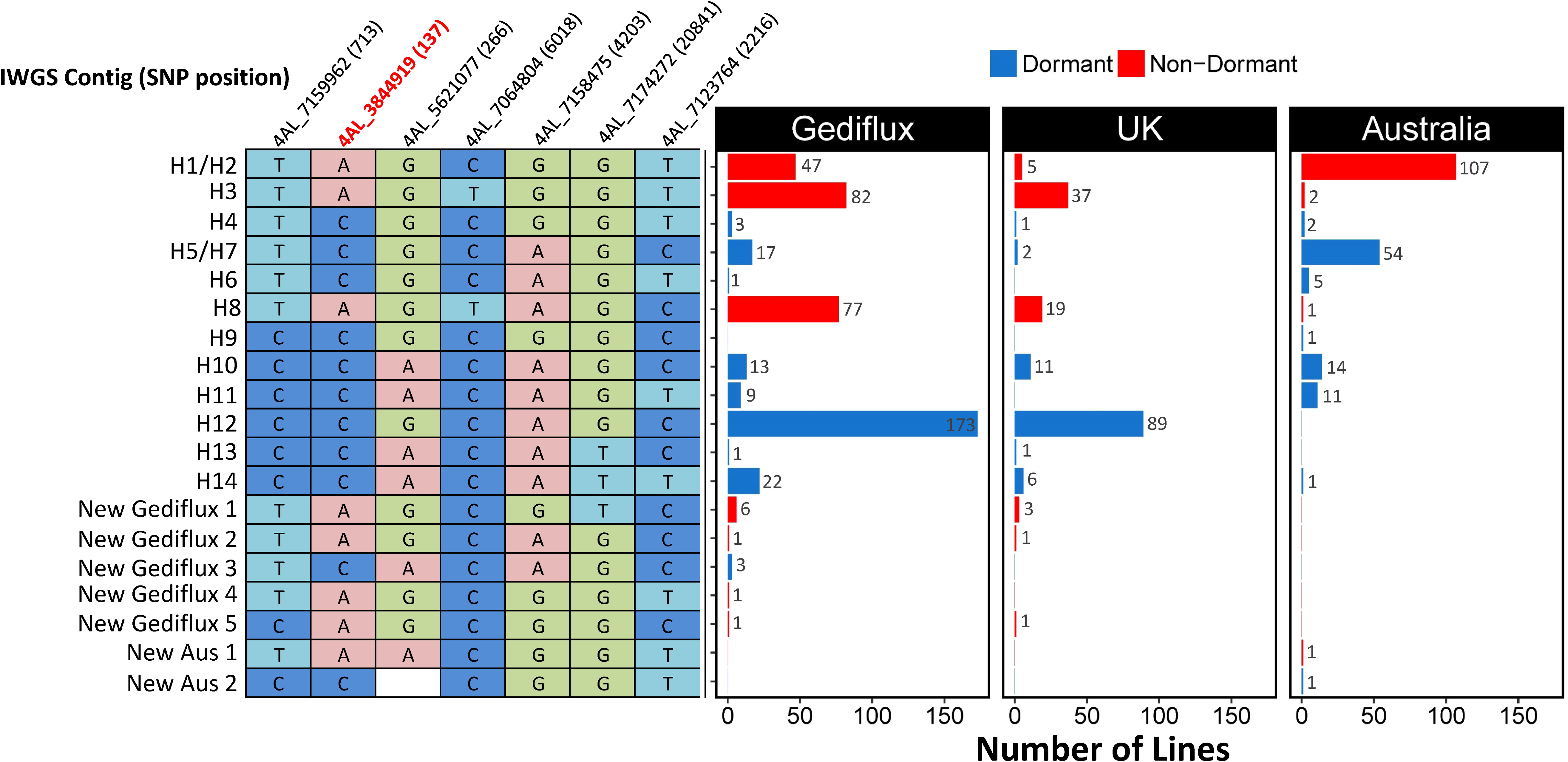
Relationship between the European, UK, Australian and HapMap *Phs-A1* Haplotypes. The distribution (bar charts) of the HapMap and unique haplotypes found in the Gediflux (European), UK and Australian germplasm using genotype information of seven of the 39 HapMap SNPs within the *Phs-A1* interval.

By combining haplotype and pedigree information for these lines we could trace, to a reasonable degree of accuracy, the founder lines for the most common resistant haplotypes in UK germplasm (Figure S1). We identified the origin of the major resistant haplotype in the UK germplasm (H12) as ViImorin-27, a French winter wheat variety released in the late 1920s (Figure 5, Figure S2). Vilmorin-27 was a direct parent and the donor of haplotype H12 for Cappelle-Desprez, a major founder variety for wheat breeding programmes in Northern France and the UK released in 1948. Haplotype H12 has since remained an important part of UK breeding programmes through varieties such as Rendezvous and Riband (released between 1985-1987).

**Figure 5:**
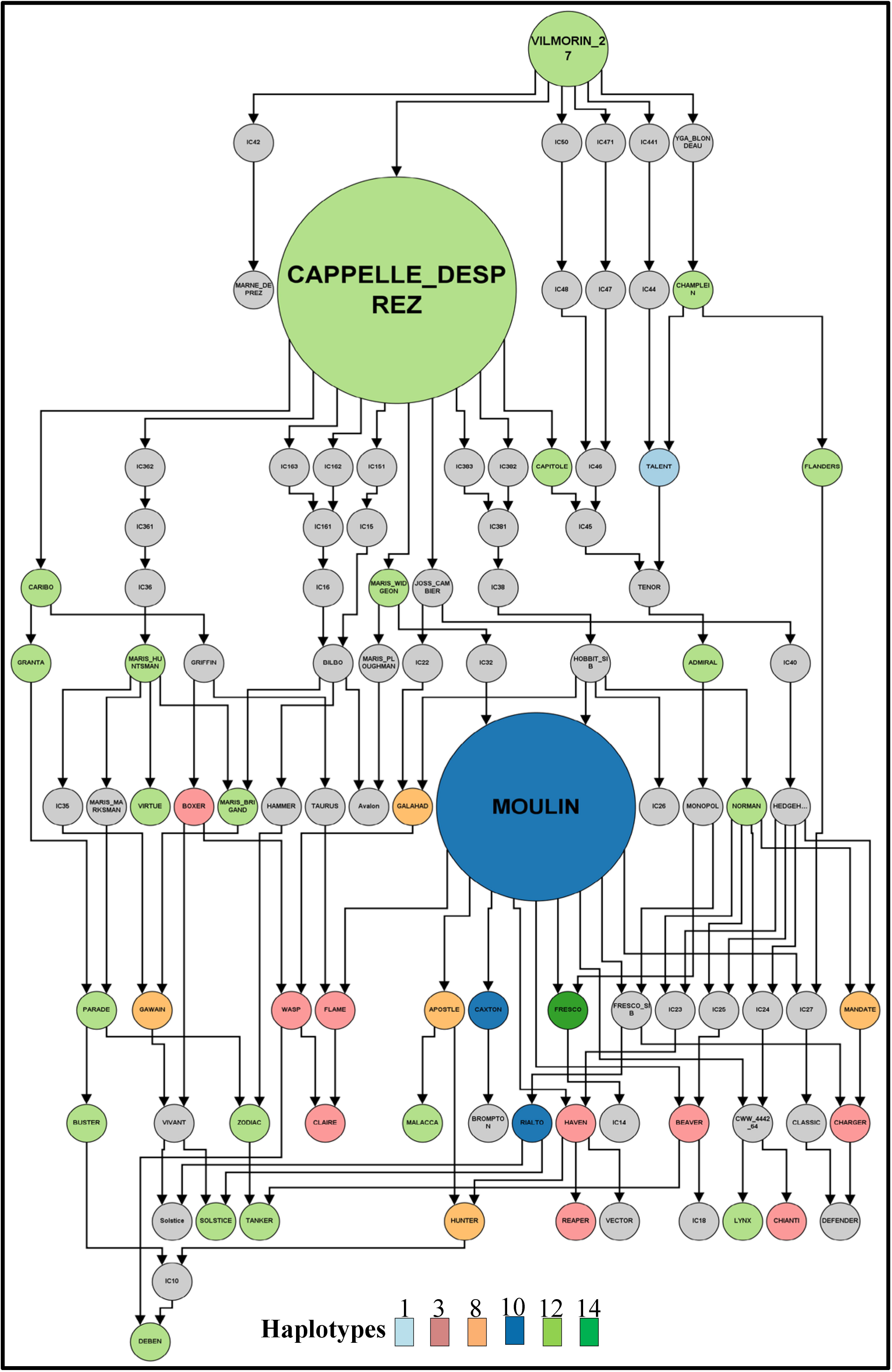
Pedigree of selected UK and European varieties highlighting the origin of the major resistant haplotype (H12). Each circle represents a variety and colours represent the different haplotypes. Nodes are size based on the number of varieties derived from the node.

Within the 200 Australian varieties we identified twelve haplotypes including ten previously identified HapMap haplotypes, and two Australian-specific haplotypes at low frequency (<1%, Table S6, Figure S3). Eight haplotypes present in 89 varieties (44.5%) have the dormant *TaMKK3-A* C allele while the other four haplotypes present in 111 varieties (55.5%) have the non-dormant *TaMKK3-A* A allele (Figure S4). This represents a near balanced distribution of both alleles in Australian germplasm. In this set, 71% of lines with the dormant *TaMKK3-A* C alleles were traced to Federation (or Purple Straw) ancestry. Across the entire set, the alternative, non-dormant allele was more associated with the presence of cv. Gabo or CIMMYT-derived material in the pedigree. These lines had a more recent average release date of 1976 compared to the lines with the dormant allele (average release date 1941).

The mean coefficient of parentage (COP) for the Australian and UK Gediflux set of lines was 0.10 and 0.11 respectively (Table 2). Within each germplasm set, the lines with the most prevalent haplotypes had higher COP values, indicating a higher degree of relatedness amongst these lines relative to the entire collection (Table 2).

**Table 2.**
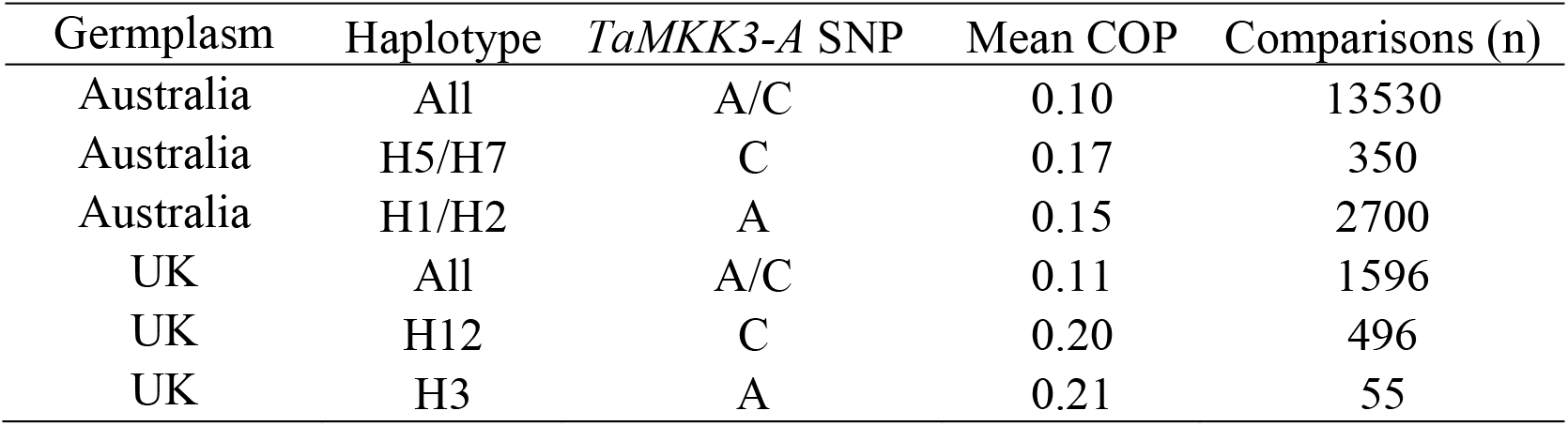
Mean Coefficient of Parentage (COP) within Australian and UK germplasm, and between groups of the most prevalent haplotypes containing dormant (C) and non-dormant (A) SNPs at *TaMKK3-A*.

| Germplasm | Haplotype | *TaMKK3-A* SNP | Mean COP | Comparisons (n) |
| --- | --- | --- | --- | --- |
| Australia | All | A/C | 0.10 | 13530 |
| Australia | H5/H7 | C | 0.17 | 350 |
| Australia | H1/H2 | A | 0.15 | 2700 |
| UK | All | A/C | 0.11 | 1596 |
| UK | H12 | C | 0.20 | 496 |
| UK | H3 | A | 0.21 | 55 |

## Discussion

### Physical Map

We characterised the *Phs-A1* interval by constructing a 1.5 Mb physical map spanning the *PM19* and *TaMKK3-A* candidate genes (Barrero et al., 2015; Torada et al., 2016) and including 16 protein-coding genes. We observed near perfect sequence and gene content conservation in the interval between hexaploid and tetraploid physical maps. A similar overall collinearity between bread wheat, barley and *Brachypodium* was also observed except for the interval between *ACC Oxidase-1* and *ERF-C* where the gene content in each species diverged (Figure S5). The *PM19* candidates where conserved across these species, whereas *TaMKK3-A* was only present in barley and wheat.

Sequence information from the BAC-based CS assembly and the whole genome shotgun Zavitan assemblies was used in a complementary manner. Neither assembly was fully contiguous across the *Phs-A1* interval, but the gaps were different in the two assemblies allowing the spanning of the complete interval. This lack of contiguity was also present in the IWGSC CS v0.4 (available at https://urgi.versailles.inra.fr/blastiwgsc/blast.php), TGAC (Clavijo et al., 2017) and Refeqv1.0 assemblies, where intervals covering *TaMKK3-A and ASC1-Like* were unanchored. While the new whole genome assemblies offer major improvements in contiguity, the available BAC physical maps will be of value to assign unanchored scaffolds or solve inconsistencies in regions were contiguity is broken.

### TaMKK3-A determines Phs-A1 effect across diverse germplasm

The 1.5 Mb physical interval which defines *Phs-A1* includes the proposed candidates *PM19* and *TaMKK3-A*, as well as other genes with potential roles in dormancy/germination regulation. For example, *ACC Oxidase-1* catalyses the last steps in the biosynthesis of ethylene – a germination promoting hormone (Matilla and Matilla-Vázquez, 2008; Linkies and Leubner-Metzger, 2012; Corbineau et al., 2014).

However, using two bi-parental mapping populations we showed linkage of *Phs-A1* to the interval between *PP1-Like – LRR Kinase 2* in UK populations, thereby excluding the *PM19* and *ACC Oxidase-1* loci as candidate genes (Shorinola et al., 2016). This was consistent with Torada et al. (2016) who identified *TaMKK3-A* as the causal gene in their mapping population and work in barley which identified the barley homologue *(MKK3)* as the causal gene for the seed dormancy QTL SD2 (Nakamura et al., 2016).

In support of this, the causal *TaMKK3-A* C660A SNP is perfectly associated with the phenotypes of 19 diverse parents of eleven mapping population in which *Phs-A1* had previously been identified. This was also the case for the parents of the MAGIC population (Yipti, Chara, Westonia, Baxter) previously used to propose the *PM19* loci as the causal gene (Barrero et al., 2015). Barrero et al (2015) proposed a promoter deletion in *PM19-A1* affecting motifs important for ABA responsiveness as the cause of non-dormancy in sprouting susceptible genotypes. The *PM19-A1* deletion and the non-dormant *TaMKK3-A* A allele are in complete linkage in all the non-dormant parents from the multiple mapping populations. However, the *PM19-A1* promoter deletion did not account for the dormant phenotype of Kitamoe, OS21-5 and SW95-50213 (Table 1). These dormant varieties have the *PM19-A1* promoter deletion associated with low dormancy, but carries the dormant *TaMKK3-A* allele. These natural recombinants suggest that *TaMKK3-A* is the causal *Phs-A1* gene. SW95-50213 is a Chinese landrace which is an important source of *Phs-A1*-mediated dormancy in Australian breeding programs. When SW95-50213 was crossed to a line carrying both *TaMKK3-A* and *PM19* dormant alleles (AUS1408), no grain dormancy QTL could be identified (Mares et al., 2005). Despite the segregation of the *PM19-A1* promoter polymorphism in this population, all lines displayed dormant to intermediate dormancy phenotype consistent with the *TaMKK3-A* genotype of their parents. Taken together, this evidence confirms the tight linkage between *TaMKK3-A, PM19*, and the *Phs-A1* phenotype, and suggest that *TaMKK3-A*, but not *PM19*, is the causal gene underlying sprouting variation associated with *Phs-A1* in diverse European, North American, Australian and Asian germplasm.

### Breeding implications

Given the identification of a number of *T. urartu* accessions with the non-dormant A allele, it is likely that the C660A mutation originates from this diploid ancestor and predates the domestication and hybridisation events that gave rise to modern bread wheat. The non-dormant allele frequency was below 15% in accessions and landraces collected previous to 1920, but rose sharply to close to 50% in more modern germplasm. It is tempting to speculate that this could be due to selective pressure by breeders over the past 70 years for the non-dormant A allele in European and Australian environments. This pressure could be driven by selection for genotypes with more rapid and uniform germination that would be associated with the non-dormant allele. However, allele frequencies for both alleles have remained overall balanced given the improved end-use quality associated with the dormant allele. This hypothesis is supported by the fact that 85% of UK bread-making varieties carry the dormant allele, compared to only 35% of feed and biscuit-making varieties.

To facilitate breeding for PHS resistance, we developed co-dominant KASP and CAPS markers for the causal *TaMKK3-A* mutation, as well as KASP markers for the wider region. We identified fourteen *Phs-A1* haplotypes in a global germplasm panel with four haplotypes for the *TaMKK3* gene itself, of which only one included the C660A non-dormant SNP. Comparison of Australian and UK haplotypes highlighted distinct frequencies in both sets with the most prevalent haplotypes containing the dormant *TaMKK3-A* allele differing in both countries. Haplotype H5/H7 is most frequent in Australian varieties, whereas haplotype H12 dominates in the UK. Interestingly, these haplotypes are either rare (<5% H5/H7 in UK) or absent (H12 not present in Australia) in the other country, suggesting distinct sources of pre-harvest sprouting resistance in Australian and UK breeding programmes. Understanding haplotypes structure across genes of agronomic interest is increasingly possible with the latest advances in wheat genomics (Clavijo et al., 2017; Uauy, 2017). It is also increasingly relevant given potential negative linkage drag associated with major phenology traits (Voss-Fels et al., 2017). The markers and knowledge generated in this study should facilitate the choice of parental genotypes for the deployment of *TaMKK3* in commercial cultivars.

The earliest line in the Australian set (Golden Drop, released 1840) carries the favourable TaMKK3-A ‘C’ SNP and also the most prevalent haplotype (H5/H7) at this locus. Golden Drop was derived from a Purple Straw/Yandilla cross and its sister line, Federation (released in 1901) become the foundation of many successful Australian cultivars due to earlier maturity and thus ability to avoid drought stress late in the growing season. Not only was Federation wheat better adapted to the Australian climate, it also had improved grain quality for milling, and so become widely adopted by breeders (Eagles et al., 2009).

The next major introduction of germplasm into Australia occurred in the 1970’s, as CIMMYT material was deployed widely by breeders seeking traits affecting height, quality and disease resistance (Brennan and Fox, 1998). Important CIMMYT parents in Australian breeding include Sonora-64, Pitic, Pavon-76, WW15 and WW80. Pedigree analysis suggests that such material could be the source of the most prevalent haplotype in Australia (H1/H2) containing the non-favourable *TaMKK3-A* allele. A high proportion of modern Australian cultivars with the non-dormant haplotype suggests opportunities may exist for the incorporation of favourable alleles at the locus.

### Future outlook

The dormant *TaMKK3-A* C allele is predominant in all the progenitor and historic germplasm evaluated in this study, suggesting that it represents the ancestral allele as proposed by Torada et al. (2016). The N220K amino acid substitution (C660A mutation) in the kinase domain results in a gain-of-function allele which reduces dormancy in wheat. This is in contrast with barley where the non-dormant *MKK3* allele is ancestral and the N260T substitution in the kinase domain results in a loss-of-function allele leading to increased seed dormancy (Nakamura et al., 2016). This provides an additional example of how for the same biological process, gain-of-function (dominant) mutations have been more readily selected in polyploid wheat compared to recessive variation in diploid barley (Borrill et al., 2015). The fact that the same gene has been selected in both species also suggests that the kinase activity of *TaMKK3-A* can be modulated to fine-tune the level of seed dormancy in temperate cereals. A better understanding of the activity and regulation of *TaMKK3-A* and its homoeologs could allow the identification of mutants (Krasileva et al., 2017) or the creating of gene edited alleles (Zong et al., 2017) with different levels of activity or the design of novel alleles with different degrees of dormancy.

## Author contributions

OS led the genotype and pedigree analysis of the UK varieties, annotation of BAC sequences, developed the KASP and CAPS marker, and analysed the HapMap data; JH performed pedigree analysis of Australian; JFGT and MJH performed genotyping of Australian varieties; MV, BB and KH constructed the 4AL physical map of CS; AD constructed the physical map of tetraploid wheat Zavitan; AT performed genotyping of Japanese varieties; JMB led the work on Australian varieties; OS, CU, JH, JMB contributed to the writing of the manuscript; OS and CU designed the experiments.

## Acknowledgements

We thank Chloe Riviere for help with plant husbandry and Dr Sergio Galvez-Rojas for advice on data visualisation. We also thank Katerina Viduka for the assistance in the genotyping, Dr Ben Trevaskis and Dr Howard Eagles for assistance in the pedigree analysis of the Australian collection, and Dr. Frank Gubler for the helpful discussions and support. We thank the International Wheat Genome Sequencing Consortium (IWGSC) for pre-publication access to the wheat genome RefSeq v0.4 and v1.0.

## Supplemental files

A single excel document includes the following supplemental tables as individual spreadsheets.

**Table S1**: BAC, Sequence and gene content information for the BAC cluster in Phs-A1 interval.

**Table S2**: TaMKK3-A Alleles of cultivars in the Watkins collection.

**Table S3**: Classification of HapMap lines according to *Phs-A1* haplotypes.

**Table S4**: KASP markers designed for haplotype analysis of Phs-A1 interval.

**Table S5**: Haplotype map of the Gediflux Collection Table S6: Haplotype map of Australian germplasm

**Figure S1**: Pedigree of UK and European varieties with their corresponding *Phs-A1* haplotype status. Each circle represents a variety and the colours represent the different haplotypes.

**Figure S2**: Pedigree of UK and European varieties with their corresponding *TaMKK3-A* allele. Each circle represents a variety and the colours represent the different haplotypes.

**Figure S3**: Pedigree of Australian varieties with their corresponding *Phs-A1* haplotype status. Each circle represents a variety and the colours represent the different haplotypes.

**Figure S4**: Pedigree of Australian varieties with their corresponding *TaMKK3-A* allele. Each circle represents a variety and the colours represent the different haplotypes.

**Figure S5**: Comparison of syntenic *Phs-A1* physical maps and contigs in *Brachypodium*, wheat and barley. Genes in the homologous intervals in *Brachypodium* (amber line), wheat (black lines) and barley (grey line) are compared against each other. Orthologous genes across genomes are joined by lines. The region of high conservation is indicated with grey background while the region of low conservation has plain background.

